# DeepPhage: leveraging the power of NGS phage display data, an in-depth analysis for the identification of Eu^3+^-binding peptides

**DOI:** 10.64898/2026.10.29.762794

**Authors:** Bloss Christoph, Huebner Amelie, Renard Clément Louis, Techert Gerda, Lederer Franziska Linda

## Abstract

**Summary:** Phage Surface Display (PSD) is a widely used molecular technique for identifying peptides and antibodies with applications in therapeutics, diagnostics, and materials science. The use of highly diverse phage libraries enables the screening of millions of ligand variants, while the integration of Next-Generation Sequencing (NGS) provides high-throughput access to the sequence composition of selected phage populations. Despite these advances, the identification of genuine target-binding ligands remains challenging due to biases introduced during library construction, biological selection and propagation, as well as the enrichment of target-unrelated peptides (TUPs). These factors distort sequence distributions and complicate the interpretation of PSD-NGS experiments, particularly when comparing multiple libraries or selection rounds. Consequently, robust computational methods are required to distinguish biologically relevant enrichments from technical artifacts. We demonstrate DeepPhage using PSD-NGS datasets from Eu^3+^ biopanning experiments, in which differences in peptide composition between experimental conditions were identified and a systematic comparison of several conditions was made possible. It was developed to address the lack of comparative analysis tools for large-scale PSD-NGS experiments by combining statistical analyses with comprehensive visualisation. These features form the basis for a more reliable interpretation of phage display selection experiments. DeepPhage provides a reproducible framework for comparative PSD-NGS analysis, which supports the identification of sequence bias, library heterogeneity and potential target-binding peptides across multiple experiments. The software is freely available as an open-source command-line application, so that it can be deployed on a high computer cluster for particularly large volumes of data.

**Availability and implementation:** DeepPhage is entirely open source and is available on GitLab https://codebase.helmholtz.cloud/phagelabs-hif/DeepPhage/-/tree/stable_release_v0.1 under the GPL v3 licence. The repository contains documentation, installation instructions, a user guide and example data. The sample data used for the demonstration is available at the online repository RODARE (https://doi.org/10.14278/rodare.4647).

## Introduction

The identification of antibodies for therapeutic and diagnostic purposes, and more recently also in the field of materials science, frequently involves the use of PSDs [1]. This is a molecular technique based on cloning of DNA fragments encoding millions of variants of specific ligands (e.g. peptides or antibodies) into the phage genome, fused to the gene encoding one of the phage’s coat proteins [2]. PSD can be used in the form of phage libraries. In this approach, random DNA is cloned into a capsid gene, resulting in the expression of peptides. The capsid protein serves as an anchor and remains structurally unchanged [3]. The gene-fusion protein is expressed and incorporated into the phage coat, so that the ligand is presented on the surface of each virus, whilst the genetic material is encapsulated within the phage particle. This physically links the presented phenotype to the genotype, making it possible to enrich specific phages simply by screening for the phenotype they display, e.g., through binding to an immobilised substrate [4]. The combination of PSD and NGS has enabled ligand identification against diverse targets, from complex biological systems to synthetic surfaces [5–7]. However, accurate binding motif discovery remains challenging due to surface heterogeneity and inherent biases introduced during library construction and phage propagation [8]. These biases, including capsid synthesis, membrane insertion, and signal peptide processing, selectively eliminate peptides disrupting phage morphogenesis while enriching sequences conferring fitness advantages [9]. Such artifacts manifest as TUPs and parasitic sequences, compromising the reliability of the experiment [10]. Sequence biases may already arise during library construction, as a key factor for highly diverse phage libraries is the random distribution of amino acids within the naïve library [11]. Deviations from uniformity lead to the over-representation, under-representation or complete exclusion of certain peptide sequences. Over-represented sequences are erroneously enriched during panning, whilst under-represented and censored sequences are prematurely eliminated from the pool, leading to the loss of potentially valuable hits. Furthermore, codon degeneracy can exacerbate this problem by leading to uneven amino acid distributions, whilst stop codons terminate translation prematurely and censor downstream sequences [12]. In addition, clones without an insert are a common construction artefact that dilutes the complexity of the library [13]. Biological bias may develop in a phage library due to constraints imposed by the host cell [10]. Another cause of sequence bias is the presence of insert sequences in the library that disrupt the protein-protein interactions required for the assembly, transport and translocation of the phage virion [14]. Ligands that are enriched not through target-specific binding but through enhanced phage fitness, known as ‘parasites’, contaminate the libraries and displace genuine binding sequences [15]. Phage display selection frequently enriches false-positive peptides lacking genuine target affinity, representing a persistent challenge in library screening. TUPs are irrelevant sequences that infiltrate target-related populations and dominate selected pools. They are categorized into selection-related (SrTUPs) and propagation-related (PrTUPs) types. SrTUPs arise from non-specific interactions with screening system components rather than the target itself. PrTUPs result from differential growth rates where faster-propagating clones outcompete slower variants. These biases are either sequence-dependent or sequence-independent. Sequence-dependent biases stem from specific amino acid motifs. Sequence-independent biases originate from mutations in phage genome regulatory regions and recombination events, enhancing pili binding, optimizing codon usage, and disrupting morphogenesis or translocation [16]. Parasitic sequences represent a PrTUP subtype with both target affinity and accelerated multiplication, offering advantages in pili binding, codon usage, and morphogenesis interference.

These biases compromise phage display accuracy by introducing off-target peptides alongside genuine binders, risking flawed conclusions. There are a number of modern databases and tools designed for the complex analysis of PSD-NGS data. In spite of that, there remains a gap in the analysis of large NGS datasets from multiple experiments, which require a comparative approach. For this purpose, we have developed DeepPhage, a standalone command-line tool that enables the comparative evaluation and visualisation of phage libraries. DeepPhage uses inserts from FASTQ files, identified by library-specific upstream and downstream markers, to extract and visualise sequence counts and their trends as part of a comparative approach. It was noted that even naïve phage library batches exhibit pronounced heterogeneity in their peptide composition and differ from one another at the amino acid, peptide and motif levels during amplification [17]. Complex biopanning experiments involving structurally more intricate target surfaces will reveal even greater variations, which is why a robust analytical framework must be in place. This multi-sample comparison allows amino acid enrichments or depletions to be observed on a position-by-position basis. It also enables pairwise comparisons using Pearson correlation coefficients, kernel density estimation and MA plots to display these differences. DeepPhage is open source and available via GitLab as a command-line tool that can be installed using a wheel file.

### Design and implementation

The functional principle of DeepPhage is presented using a dataset generated from several Ph.D.™-12 phage library experiments involving biopanning against Eu^3+^ ions immobilised on nitrilotriacetic acid (NTA) and coupled to agarose beads (PureCube 100 NTA Agarose, Cube Biotech, Monheim, Germany) [18]. DeepPhage currently consists of three modules: (i) processing, generates peptide sequence counts and some basic plots for data overview, (ii) statistical evaluation of the frequency of peptide sequences, normalisation and/or transformation of data, calculation of amino acid enrichment or depletion, determination of statistical parameters using tests, and visualisation of paired sample data, and (iii) data clustering, involving quality control steps, data exploration and trimming, parameter setting as well as model fitting. DeepPhage requires FASTQ files from PSD experiments as input. In addition, the peptide sequence count data must be collected in a directory and named according to a documented specified pattern.

### Data preparation

We have received the raw sequence data generated by the Illumina NextSeq system in paired-end read format. The FastQC toolkit was used to assess the quality of the reads [19]. The reads were merged using BBMerge of the BBMap package and reads with a phred score below 30 have been trimmed [20, 21]. Both steps were combined in a home-brew bash script.

### Processing

The processing module is the entry point in the workflow; it processes FASTQ files and extracts library-specific inserts. In order to use this software, users need to specify certain information. There are three mandatory inputs: (i) infile, the path to the file containing the merged reads in FASTQ format, or, if the file is large, in compressed .gz format; (ii) out, the path to a user-specified output directory; and (iii) lib, the type of phage library used during a PSD experiment enabling the identification of nucleotide inserts. Found inserts, translated insert sequences and counts of translated insert sequences are each stored to an individual .csv file. Optional parameters allow for the generation of a quantile bar chart of the translated insert sequences (see Fig. S1.1), a heatmap showing the amino acid enrichment per position and an information logo (see Fig. S1.2) illustrating the information content of the amino acids per position.

The module begins by defining global paths, including root directory, temporary directory for pickle and all output paths. The isolation of nucleotide inserts involves screening for randomised inserts from merged sequences of displayed peptides. When working with commercially available phage libraries, each insert, regardless of gene region, is flanked by markers located ‘upstream’ and ‘downstream’ of the region. The phage libraries are bound to specific marker constructs [22]. A fixed selection of standard phage libraries is available. Currently, the Ph.D.™-7, Ph.D.™-12, Ph.D.™-C7C, MIM™-C10, f88-Cys4, X15, and LX4 are supported. Using a generator function, the merged FASTQ file is read in. This removing the header, the separator and the quality line. The lib argument is used to specify the relevant markers and insertion lengths. The nucleotide sequences are retrieved via regular expression. The result is a generator object containing only nucleotide inserts, which is converted into a vectorised list. This list is used to translate nucleotide inserts to peptide insert sequences using a NNK modified version of the Bacterial, Archaeal and Plant Plastid Code (transl table=11) compiled by Andrzej (Anjay) Elzanowski and Jim Ostell at National Center for Biotechnology Information (NCBI), Bethesda, Maryland, U.S.A. https://www.ncbi.nlm.nih.gov/Taxonomy/Utils/wprintgc.cgi (accessed on 11 Mai 2026). To ensure quality, NNK translation is checked against transl table=11. If NNK translation is not possible due to ambiguous codons, it is replaced by the native translation. Peptide insert sequences are grouped according to unique sequences, counted and stored, and their fraction of the total number of peptide sequences is calculated.

For visualisation three options are given. By calculating the frequencies of individual peptide sequences and the overall frequencies, users can plot five-percentile quantiles as bars, resulting in 20 bars summarizing count accumulation. The heatmap can be generated using a position-specific scoring matrix (PSSM) for amino acid residues, with pseudo counts and the background frequencies derived from the NNK codon table. Using the same matrix the information content of the amino acid residues at each position can be calculated, in order to generate a global sequence logo. Additional information from the processing module can be found in the Appendix S1. In a nutshell, the file containing the count values for the unique peptide sequences is particularly important as a basis for the next module.

### Statistical evaluation

The statistical analysis module is used to evaluate previously determined counts of peptide insert sequences. The focus is particularly on the intersection and the exclusion of relative samples. The following four parameters are mandatory and must be used in this context: (i) sampledir, the directory for count files of samples. File names need to follow a specific naming convention, i.e. ‘XYZ SampleN countsM.csv’. Where *N*, is the sample number e.g. biological replicate and *M* is the number of the technical replicate. (ii) lib, the single, pre-determined phage library type (iii) normalisation, supported normalisation/transformation methods, (iv) non-parametric, offers a choice of non-parametric tests. To run these tests, two additional parameters, ‘amount’ and ‘frequency’, are necessary. ‘Amount’ changes the number simulated peptides. ‘Frequency’ refers to an amino acid distribution based on codon usage. Additionally, the output is saved in the ‘out’ directory within the programme directory, but changing the ‘outdir’ parameter enables selection of a different output path. Further options include options include generating Quantile-Quantile (Q-Q) plots for visualising sample normality, performing D’Agostino’s K^2^ test and Anderson-Darling test, plotting position-specific amino acid charts, and a combination plot with MA plots, Kernel Density Estimation (KDE) plots, and sample correlations.

The statistics module first sets up global paths for the program, including the root directory, the pickle directory, and the input and output paths for statistical information. It then parses the files containing the peptide sequence counts from the input directory and creates a data frame. This data frame has *n* rows, where *n* is the number of sequences. With *p* + 1 columns in the data frame, where *p* is the number of samples. Sample names are relabelled based on the sample name, sample number *N* and replicate number *M*. The index holds the peptide sequences and the values of the frame correspond to the counts of each sequence per sample. Zero values are replaced by the appropriate proportion of the ratio of the row total and the sum of the row (including zero values), while the resulting row total remains unchanged. In order to determine whether a dataset follows a normal distribution, the quantiles of the observed data are compared with the theoretical quantiles of a normal distribution. This is used to create a Q-Q plot, see. In order to observe the position-specific enrichment and/or depletion of amino acids within the insert, a position-specific bar chart can be generated. This can be used when analysing a biopanning run involving multiple samples per biopanning round. The transformed data frame is expanded using the numerical values in each column, and a nested dictionary is created, containing one data frame per column. Each column in this dictionary of data frames corresponds to a sample. At first, for each data frame in the dictionary, a position count matrix (PCM) is build. For speed-up and matrix building it converts string sequences of alphabet:

‘ACDEFGHILKMNPQRSTVWY*’ to integers. Therefore, stacking integer sequences (*seq*) into a 2D-array of shape (n_*seq*_, len_*seq*_). All positions are initialized with a pseudo count of one and occurrences of encoded letters per column are counted. Next, the PCMs are each transformed to position frequency matrix (PFM) with column wise normalisation. This is done by calculating the total values for each position per column and adding the totals in each column. The result is then divided by the corresponding column total for each row. Alongside, a dictionary with data frames for background distributions is created. This is used to transform the PFM to a PSMM with a normalized background for amino acids on specific position. The amino acid background frequency for position *i* was sourced from the preceding data frame of the background dictionary *d*_*b*_ with the same position (*d*_*b*_*−* 1[*i*]). For the first PSSM the background is derived from the corresponding data frame itself. In order to visualize the results, a colour map is created based on all unique sample names, showing the final PSSM scores for each sample. A visualisation can be seen in Fig. S2.2.

Normalisation/transformation is done with one of the selected methods. There are simple methods such as Counts per Million (CPM), Trimmed Mean of M-Values (TMM) [23] and Median of Ratio Normalisation (MRN) [24]. To calculate summary statistics, particularly for analysing skewness and kurtosis, users can employ the built-in D’Agostino’s K^2^ test. The Anderson-Darling test is intended for data that follows a specific distribution. The test examines the null hypothesis that a sample is drawn from a population that follows a specific distribution. The default setting is ‘norm’ for the normal distribution. For visualization purposes of these normalised or transformed values, a plot matrix showing pairwise comparisons of the samples is generated. Above the diagonal, the pairwise correlation between the samples is displayed, calculated using the Pearson correlation coefficient. Along the diagonal, KDE plots for individual samples are shown. Below the diagonal, MA plots are displayed in pairs; where applicable, the normalisation factor is also shown. Supplementary information about the pairwise matrix plot can be seen in Fig. S2.3.

To perform a non-parametric tests random *in-silico* peptides are generated. This is based on the ‘Random Amino Acid and Nucleotide Sequence Generator’ [25], which forms part of the Bactome package and has been adapted to handle peptides and permutations while ensuring efficient computation times. The amount is set to 1000 by default, but can be adjusted. The length is fixed and determined by the library used. There are three ways to specify the frequencies of amino acids, two of them involve user-defined sources. User data can be provided as a .txt file containing a frequency table for amino acids, or as a .csv file, with the frequencies calculated from the amino acid read counts. When using a .csv file, specify the column in the peptide count file to define the amino acid distribution. This can be kept at the default setting if created by DeepPhage processing module. This generated artificial data set is compared with the real data set, using Mann-Whitney *U* rank test, Wilcoxon signed-rank test, Friedman test and a chi-squared test of independence of variables in a contingency table [26]. Each test is carried out using 10 000 permutations, with the exception of the Friedman test, for which no permutations are carried out. For further information about the statistical evaluation see Appendix S2.

## Results

To demonstrate DeepPhage, we used paired-end sequencing data from four experiments, all involving Ph.D.™-12 phage libraries. The first experiment performed a biopanning using Eu^3+^ ions immobilised on NTA and coupled to agarose beads, while the second biopanning was conducted without immobilized Eu^3+^ ions as a control. Both were eluted biologically using *E. coli* cells. The last two experiments follow the same design, with the difference that elution was carried out chemically with glycine-HCl. [18]. The original goal of this study was to identify Eu^3+^-binding peptides using PSD. The study was designed so that enriched sequences across different elutions could be compared and correlated. The data was quality controlled using FastQC version 0.12.1, paired-end sequences were created using BBMerge from BBMap version 39.26 and preprocessed following the steps described in the data preparation section. Parts of the output from the processing phase and the statistical analysis is shown in Fig 1. Further results relating to data processing can be found in Appendix S1, whilst further results relating to the statistical analysis can be found in Appendix S2. The global position-specific amino acid enrichment of the insert sequences, which were isolated by biopanning using pure NTA-agarose beads as a control and chemical elution is displayed in Fig 1(A). It shows a strong depletion of proline at position one, as well as a consistent depletion of cysteine and the stop codon across all positions. It is also evident that arginine is depleted. Interestingly, the depletion decreases across all positions. Lysine is noticeably reduced at positions one and two, while leucine is reduced only at position one. The most clearly enriched amino acids are asparagine from positions one onwards, with a decreasing value thereafter, as well as histidine, which is also particularly enriched at position one and then decreases, aspartic acid shows a similar behaviour, and alanine, albeit only at position one. By comparison, the heatmap in Fig 1(B) shows biopanning using Eu^3+^ ions and chemical elution. It presents a similar pattern, with a uniform depletion of cysteine and stop codons at all positions, as well as a distinct depletion of proline at position one. The decreasing depletion of arginine across all positions is also evident. Lysine shows a slight enrichment at position one, but no enrichment at position two. Leucine remains depleted at position one, but shows a slight enrichment at position nine. However, it also shows a marked enrichment of histidine at positions two to five, as well as at positions ten and eleven. In contrast to the control, asparagine is heavily enriched at positions six and seven and is relatively depleted at all other positions. Glutamine is heavily enriched at position ten, as are proline at position eleven and phenylalanine at position twelve. The paired plot with CPM normalised insert counts, in Fig 1(C), shows the Pearson product-moment correlation coefficients in the upper right. It indicates a slight correlation between the control (Sample1) and target samples (Sample2) for biological elution, and between the control sample for biological elution and the control sample for chemical elution (Sample3). The diagonal shows the KDE plots for each sample. It can be seen that the probability distributions for Sample1 to Sample3 are pretty similar. The plot for Sample4, on the other hand, has a different shape, with two curves that are clearly distinct from one another. The MA graphs are shown at the bottom left. The plot for the pair of Sample1 and Sample2 shows that there appear to be some inserts that are differentially expressed, with a *log*_2_*FC*~0. Furthermore, it can be seen that, in the range of lower expression levels, there are condition-specific inserts. It appears that, in the remaining MA plots, there is a biological expression trend, with very few insert variants showing differing expression levels. As a control, the paired plot was generated using TMM normalisation, as shown in Fig 1(D). The Pearson product-moment correlation coefficients show no change. In the KDE plots, the mean values have changed overall, but the curves of the individual probability distributions look similar. The main difference is clearly recognizable in the MA plots. The biologically eluted samples, paired with the chemically eluted control sample, behave similarly to the plots from the CPM normalisation. The key difference becomes clear when the data is paired with that from Sample4. This shows that the inserts in Sample4 are upregulated (*log*_2_*FC >* 0) in all other samples. This is also evident from the normalisation factor line of *log*^2^*mean*, which lies between 8.01 and 8.37 for these samples. Further varied results relating to other normalisation methods can be viewed in Appendix Sample2, figure xyz. Nevertheless, it is vital to mention that the diversity and especially the signal-to-noise ratio can explode in PSD experiments [16]. Given the complex compositional nature of this data, it is recommended that all available analytical methods be utilised. DeepPhage offers a wide range of statistical and visual methods for evaluating PSD experiments. This enables the relevant experimental conditions to be compared quickly and directly as part of a thorough analysis.

**Fig 1.**
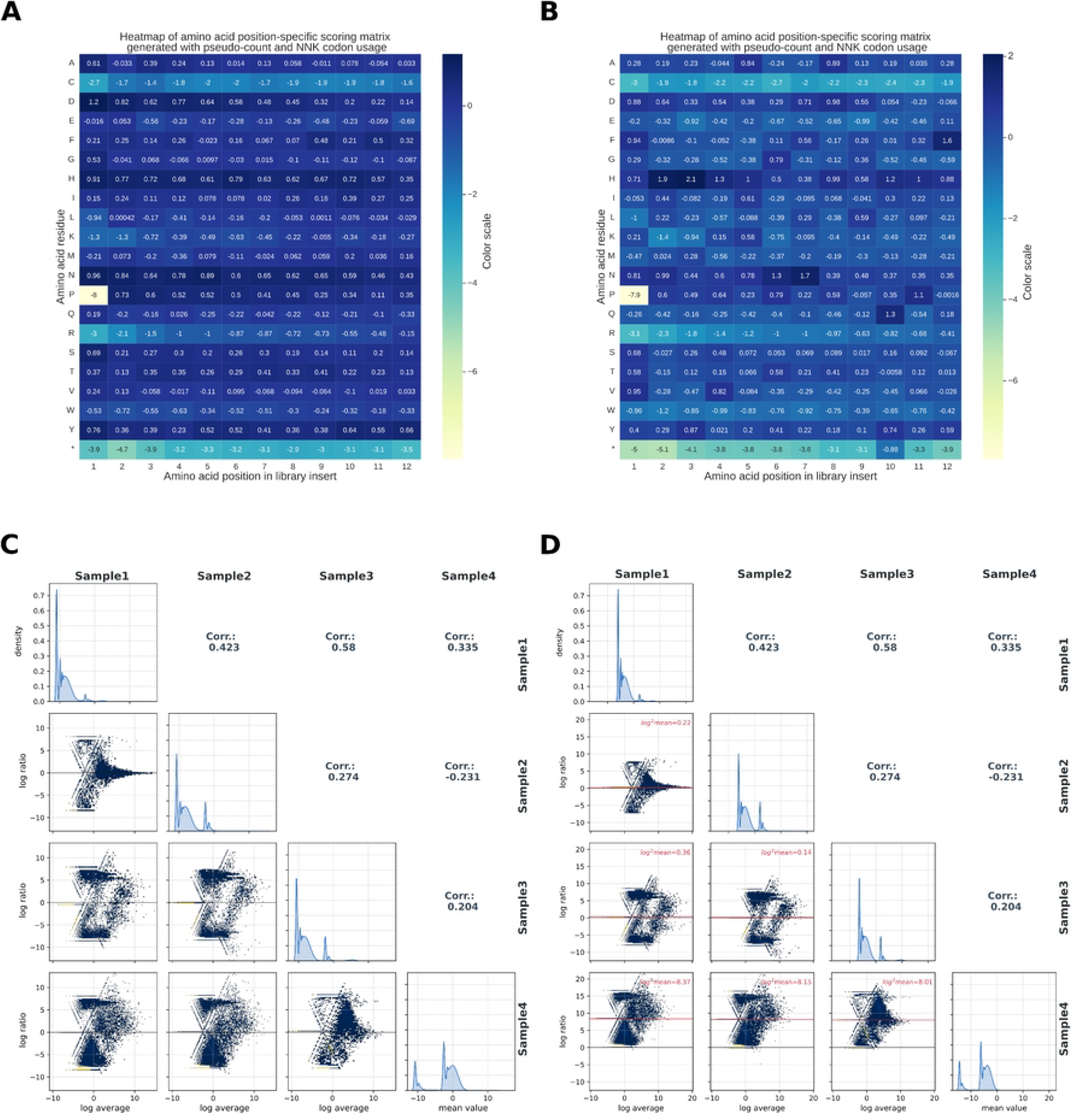
Analysis and visualisations of two PSD experiments to identify Eu^3+^-binding peptides using DeepPhage. (A) PSSM of the amino acids in the 12-mer insert from a biopanning assay against pure NTA-agarose beads and chemical elution, presented as a heatmap. The positions shown below are the amino acids on the left, and the colour map indicates an enrichment (dark) or depletion (light) compared with the expected value, which is calculated from the pseudo-count and background probability. (B) heatmap of the biopanning with immobilised Eu^3+^-ions. A comparison of the two heatmaps can reveal target-specific changes. (C) A pairwise matrix plot of the CPM-normalised count values, showing correlations, KDE plots and MA plots. The yellow dots in the point clouds indicate that points are overlapping. It is noticeable that both the two biologically eluted samples (Sample1 and Sample2) share a larger number of inserts and the chemically eluted control sample with the biologically eluted control sample (Sample1 and Sample3). (D) As a control the pairwise-matrix plot was created with TMM normalisation. The key difference becomes apparent when the data are compared with Sample4: this reveals a significant upregulation (*log*_2_*FC >* 0) of inserts in all other samples. This is also evident from the normalisation factor (red line), which is heavily skewed towards all other samples.

## Discussion and conclusion

The results generated using phage display technology may often contain TUPs and true binding peptides, which can lead to potential errors, misleading results and incorrect conclusions. It is therefore important to validate biopanning data by identifying and distinguishing between them [27]. There are databases such as BDB [28] or TUPDB [29] that can be used, as they are still being maintained. A number of standalone programmes or scripts have also been developed to support the analysis, but these are usually limited to specific use cases. SAROTUP [30], PHASTpep [22], PuLSE [27] and PhageSeqAnalysis [7]. However, these tools are designed for small-scale assessment that do not allow for a direct, statistical comparison of different experimental conditions (e.g. control versus target or the course of a biopanning experiment involving several rounds of panning). DeepPhage fills this gap and offers various ways to access the data and different perspectives on it. The methods presented allow for more comprehensive analyses and comparisons across multiple experiments. When combined with statistical analysis and visualisation of DeepPhage, they can deliver initial indications for distinguishing between peptides with genuine binding and false-positive peptides. It can be used for any type of experimental setup, whether for analysing ongoing biopanning rounds or for comparing multiple target and control replicates. The input options can be customised to a large extent, but must be provided together with the library used in the experiment. The tool made it possible to detect a direct difference in the amino acid composition of the two chemically eluted samples. Furthermore, it was possible to compare all samples simultaneously using the pairwise plot matrix. This made it possible to compare two samples at a time by visualising the relationship between the magnitude of the measured values and the differences between them under the various conditions. For further statistical analysis, the samples must comply with a strict naming convention. Statistical tests, as well as normalisation and transformation, can be specified individually, as can parameters for permutations and the background distribution of individual amino acids. Position-specific enrichment statistics, visualisations of pairwise comparisons and non-parametric tests can be used to address issues with the signal-to-noise ratio and to identify differences between two populations. By addressing the gap in the large-scale analysis of PSD-NGS data from multiple runs or experiments, DeepPhage facilitates a better understanding of the underlying mechanisms and the formulation of hypotheses. Still, we could go much further. Recent years have shown that unsupervised machine learning can be used to gain even more insights and scan for different subgroups in experimental data [31]. We think this is an opportunity to test ensemble clustering methods and gain a better understanding of the difference between true and false binders. This enables clustering across a wide range of distance metrics, transformations and validation metrics that take into account high levels of noise and variance [32], making it ideal for the noise-to-signal ratio problem.

## Availability and future directions

DeepPhage is fully open source (under the GPL v3 licence) and available on GitLab https://codebase.helmholtz.cloud/phagelabs-hif/DeepPhage/-/tree/stable_release_v0.1. It requires no specialised hardware. Software dependencies other than standard Python packages are set up automatically, and it runs on standard computers. The repository includes full documentation, installation instructions and a user guide which uses example data to present typical use cases similar to those described in the manuscript. The repository contains a smaller example datasets for testing the functionality of the package.

As PSD-NGS datasets are plagued by signal-to-noise distortions, our next step will be to apply clustering methods that enable us to distinguish between signal and noise in these highly variable datasets and thus extract TUPs. One inspiring piece of work in this regard is from Sloth et al. 2022 [31]. On this basis, it would be possible to significantly refine the analysis and identification of binding ligands.

## Data availability

The sample data used in this study is available on the data repository RODARE (https://doi.org/10.14278/rodare.4647).

## Supporting information

**Fig. S1.1 Quantile bar plots**.

**Fig. S1.2 Sequence logos**.

**Fig. S2.1 Quantile-Quantile plots**.

**Fig. S2.2 Per position amino acids enrichment plot**.

**Fig. S2.3 MRN pairwise matrix plot**.

**Appendix S1 Further method explanation and show-casing for the processing module**.

**Appendix S2 Further method explanation and show-casing for the statistical evaluation method**.

## Acknowledgments

The authors would like to thank N.Schoenberger and S.D. Harter for their support and assistance in refining the wording and for their advice on graphical details. We would like to thank H. Hintersatz for proofreading and formulation suggestions.

